# Annotating high-impact 5’untranslated region variants with the UTRannotator

**DOI:** 10.1101/2020.06.03.132266

**Authors:** Xiaolei Zhang, Matthew Wakeling, James Ware, Nicola Whiffin

**Author notes:** Contributed equally (James Ware, Nicola Whiffin).

## Abstract

**Summary:** Current tools to annotate the predicted effect of genetic variants are heavily biased towards protein-coding sequence. Variants outside of these regions may have a large impact on protein expression and/or structure and can lead to disease, but this effect can be challenging to predict. Consequently, these variants are poorly annotated using standard tools. We have developed a plugin to the Ensembl Variant Effect Predictor, the UTRannotator, that annotates variants in 5’untranslated regions (5’UTR) that create or disrupt upstream open reading frames (uORFs). We investigate the utility of this tool using the ClinVar database, providing an annotation for 30.8% of all 5’UTR (likely) pathogenic variants, and highlighting 31 variants of uncertain significance as candidates for further follow-up. We will continue to update the UTR annotator as we gain new knowledge on the impact of variants in UTRs.

**Availability and implementation:** UTRannotator is freely available on Github: https://github.com/ImperialCardioGenetics/UTRannotator

**Supplementary information:** Supplementary data are available at bioRxiv.

## Introduction

Upstream open reading frames (uORFs) are short sequences within 5’UTRs that regulate the rate at which the downstream coding sequence is translated into protein. Variants that create or disrupt uORFs (uORF-perturbing variants) have been shown to cause rare disease (Calvo, Pagliarini and Mootha, 2009; Whiffin *et al.*, 2020). We recently used data from the Genome Aggregation Database (gnomAD) to systematically characterise the deleteriousness of different categories of uORF-perturbing variants and prioritise those that are more likely to be disease causing (Whiffin *et al.*, 2020). Current variant annotation approaches focus on the impact of protein-coding variants, with only limited annotation of predicted consequences for non-coding variants. For example, the Ensembl Variant Effect Predictor (VEP) (McLaren et al. 2016), only annotates variants within UTRs as 3’ or 5’ to the coding sequence, without any further information about their predicted effect.

To aid the assessment of high-impact uORF-perturbing variants, we have developed a plugin for VEP to identify 5’UTR variants that create upstream start sites (uAUGs), disrupt the start or stop codon of existing uORFs, or shift the frame of an existing uORF. In each case, the tool outputs detailed annotations that allow the user to predict the likely impact of the variant on protein translation.

Recently, the MORFEE tool was described (Aïssi *et al.*, 2020), however, it is limited to annotating single nucleotide variants (SNVs) that create uAUGs. The UTRannotator is, to our knowledge, the first comprehensive annotation tool for 5’UTR uORF creating and disrupting variants. Our tool has initially been created to characterise the impact of uORF-perturbing variants, however, it will be updated to annotate additional UTR variants as we learn how to interpret these for a role in human disease.

### Approach

For any SNV, 1-5bp small insertion/deletion (indel) or multi-nucleotide variant (MNV) in a 5’ UTR, we first summarize the number of uORFs in the 5’UTR in the reference sequence. Then, for each variant within the 5’UTR we evaluate whether it would have any of the following consequences, on any annotated transcript: (1) creating a new start codon AUG to introduce a new uORF; (2) removing an existing start codon AUG; (3) removing the STOP codon of an existing uORF; (4) disrupting an existing uORF with a frameshift deletion or insertion, whose number of nucleotides inserted or deleted is not a multiple of three.

To enable evaluation of the effect of each variant, the UTRannotator outputs detailed annotations for each type of uORF-perturbing variant (**Table 1**). This includes describing the subtype of uORF created and/or disrupted (i.e. whether this is a distinct uORF with a stop codon in the 5’UTR, or an ORF that overlaps the coding sequence either in-or out-of-frame), and the strength of the created and/or disrupted uORF start site match to the Kozak consensus sequence (Kozak 1989). For a variant disrupting an uORF, we also evaluate whether the uORF has any experimental evidence of translation, by assessing a curated list of uORFs previously identified with ribosome profiling from the online repository of small ORFs (www.sorfs.org) (Olexiouk, Van Criekinge and Menschaert, 2018). Users can also use their own customised list of translated uORFs.

Since a 5’UTR can have multiple existing uORFs, for each 5’UTR variant we output the annotations for all disrupted uORFs.

The time complexity of our implementation is linear to the number of input variants. The ratio of running time without the plugin to that with the plugin, tested on 1000 random variants (60% annotated as 5’UTR variants) is 1.02-1.07 (5 replications).

**Table 1.** Details of the annotations provided for different categories of uORF-perturbing variants.

| Consequence | number of existing uORFs | KozakContext: sequence and strength | Start distance to CDS | Start distance to STOP | With translation evidence | uORF subtype | Other annotations |
| --- | --- | --- | --- | --- | --- | --- | --- |
| uAUG_gained | ✓ | ✓ | ✓ | ✓ |  | ✓ | Start distance from cap |
| uAUG_lost | ✓ | ✓ | ✓ | ✓ | ✓ | ✓ |  |
| uSTOP_lost | ✓ | ✓ |  |  | ✓ |  | Whether there is an alternative STOP, alternative stop distance to CDS, frame of disrupted uORF with CDS |
| uFrameshift | ✓ | ✓ | ✓ |  | ✓ | ✓ (ref and alt) |  |

## Results

To show the utility of our UTR annotator tool, we annotated all 5’UTR variants interpreted as pathogenic/likely pathogenic and uncertain significance from ClinVar (version 202005) (Landrum *et al.*, 2018). These variants do not have a coding annotation on any transcript. However, we note that 5’UTR variants are under-represented in ClinVar as they are rarely sequenced and/or reported.

There are 97 Pathogenic/Likely pathogenic 5’UTR variants in ClinVar (97/113,969=0.085% of all ClinVar Pathogenic/Likely pathogenic). 91 are 1-5bp small variations, 28 of which (30.8%) are annotated as creating or disrupting uORFs by our plugin (**Figure 1; Supplementary Table 1**). We examined the evidence behind the reported clinical significance for each variant, and found 14 (50%) have previously been attributed to a uORF-perturbing mechanism.

**Figure 1.**
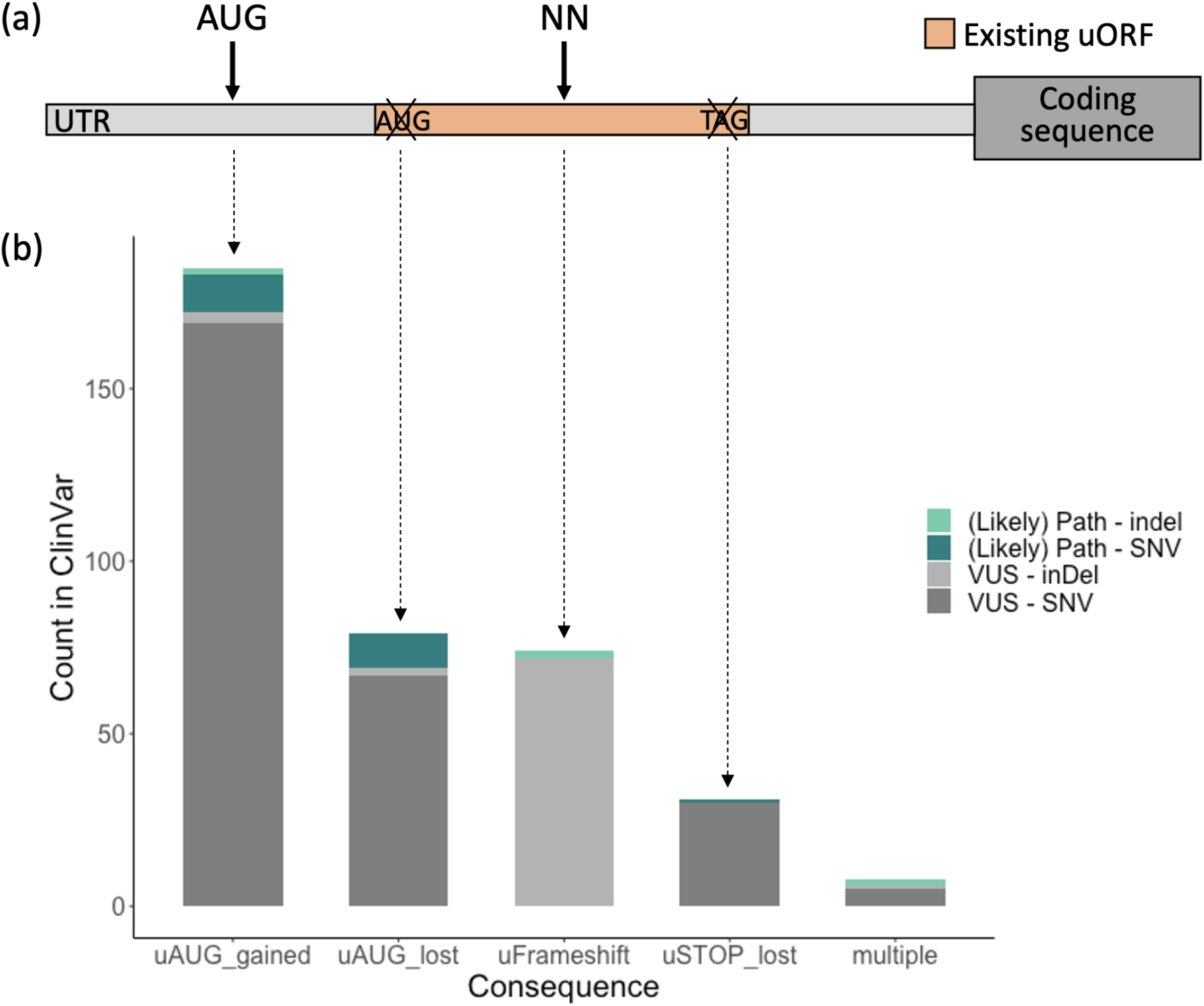
5’UTR variants in ClinVar annotated by the UTRannotator. (a) A schematic showing the four distinct consequences of 5’UTR variants annotated by the tool: those that create an upstream AUG (uAUG_gained), those that disrupt the start or stop sites of existing upstream open reading frames (uORFs; uAUG_lost and uSTOP_lost respectively) and those that cause a frameshift in the sequence of the uORF (uFrameShift). (b) The counts of each variant category that are classified as Pathogenic/Likely Pathogenic (teal) or Uncertain Significance (VUS; grey) in ClinVar.

There are 5,128 5’UTR variants of uncertain significance (VUS) reported in ClinVar (5,128/255,691=2% of all VUS), 4,966 of which are 1-5 bp small variations. Our plugin annotated 349 of these (7.0%) as creating or disrupting uORFs, on at least one annotated transcript (**Supplementary Table 2**).

We used the detailed annotations from the UTRannotator to illustrate how to prioritise 5’ UTR VUS that are most promising for further follow-up. We first restricted to variants that form new overlapping ORFs (oORFs) with start sites that are Strong or Moderate matches to the Kozak consensus sequence, or that are uORFs with documented evidence of translation, as we previously showed that variants with these consequences are under strongest negative selection (Whiffin *et al.*, 2020). Finally, we took variants in genes previously identified as having a ‘High’ likelihood that uORF-perturbation could be an important disease mechanism (Whiffin *et al.*, 2020). Through this approach, we identified 31 potential ‘high-impact’ ClinVar 5’UTR VUS (**Supplementary Table 3**).

## Discussion

We have created a freely available tool, as a plugin to the Ensembl VEP, that annotates variants that create or disrupt uORFs. The output from the tool can be used to predict the possible impact of variants identified in patients for a role in disease. It is also directly applicable to annotate 5’UTR variants from other eukaryotes.

We note several limitations to our tool. Firstly, the UTRannotator has been configured to annotate only variants up to 5bps in length. We included this length restriction for two reasons: (1) the annotation of longer indels is tricky, as the chance of variants having multiple possible annotations is increased, and (2) the impact of larger indels that add or remove large stretches of UTR is currently unknown. We also currently only consider uORFs with canonical AUG start sites. It is known that many translated uORFs use non-canonical start sites (McGillivray *et al.*, 2018). More research is needed into the impact of variants that create or disrupt these non-canonical uORFs in human disease.

For the initial tool release, we have included four variant types that create or disrupt uORFs, however, we will continue to develop the UTRannotator to include additional types of UTR variants.

